# ADPKD-Causing Missense Variants in Polycystin-1 Disrupt Cell Surface Localization or Polycystin Channel Function

**DOI:** 10.1101/2023.12.04.570035

**Authors:** Kotdaji Ha, Gabriel B. Loeb, Meyeon Park, Mohona Gupta, Yukako Akiyama, Jillian Argiris, Aide Pinedo, Christine Haewon Park, Nadav Brandes, F. Ritu, Chun Jimmie Ye, Jeremy F. Reiter, Markus Delling

**Affiliations:** Department of Physiology, University of California, San Francisco, San Francisco, CA, USA; Division of Nephrology, Department of Medicine, University of California, San Francisco, San Francisco, CA, USA; Cardiovascular Research Institute, University of California, San Francisco, San Francisco, CA, USA; Division of Rheumatology, Department of Medicine, University of California, San Francisco, San Francisco, CA, USA; Division of Rheumatology, Department of Medicine, Bakar Computation Health Sciences Institute, Parker Institute for Cancer Immunotherapy, Gladstone-UCSF Institute of Genomic Immunology, Institute for Human Genetics, Department of Epidemiology & Biostatistics, Department of Bioengineering & Therapeutic Sciences, University of California, San Francisco, San Francisco, CA, USA; Department of Biochemistry and Biophysics, Cardiovascular Research Institute, University of California, San Francisco, San Francisco, CA, USA; Chan Zuckerberg Biohub, San Francisco, CA, USA

## Abstract

Autosomal dominant polycystic kidney disease (ADPKD) is the leading monogenic cause of kidney failure and affects millions of people worldwide. Despite the prevalence of this monogenic disorder, our limited mechanistic understanding of ADPKD has hindered therapeutic development. Here, we successfully developed bioassays that functionally classify missense variants in polycystin-1 (PC1). Strikingly, ADPKD pathogenic missense variants cluster into two major categories: 1) those that disrupt polycystin cell surface localization or 2) those that attenuate polycystin ion channel activity. We found that polycystin channels with defective surface localization could be rescued with a small molecule. We propose that small-molecule-based strategies to improve polycystin cell surface localization and channel function will be effective therapies for ADPKD patients.

---

Autosomal dominant polycystic kidney disease (ADPKD) is a common monogenic disease in humans and accounts for ∼10% of global kidney failure^1^. Because the molecular mechanisms underlying ADPKD remain largely unknown, treatment remains extremely limited. ADPKD is characterized by significant allelic heterogeneity with over 1200 pathogenic or likely pathogenic variants identified^2^, with most pathogenic variants restricted to individual families, even in large cohorts^3,4^. Most ADPKD (80%) results from heterozygous variants in polycystin-1 (PC1), most remaining cases (15%) are caused by heterozygous polycystin-2 (PC2) variants^3,5,6^. PC1 and PC2 can assemble into a heteromeric TRP channel with 1:3 stoichiometry (hereafter called the polycystin complex). The polycystin complex localizes to the primary cilia of the kidney tubular epithelium, where it is thought to exert its physiological function^1,7–9^.

Interestingly, a significant proportion of ADPKD arises from nontruncating, primarily missense, variants in PC1 and PC2^3,4,10^. However, the absence of functional assays to classify such variants has remained a major challenge in ascertaining their pathogenicity. This lack of information is in stark contrast to the current understanding of another life-threatening monogenic disease, cystic fibrosis (CF). CF results from loss-of-function variants in the cystic fibrosis transmembrane conductance regulator (CFTR) ion channel^11^. Pathogenic variants in CFTR either disrupt cell surface localization, CFTR channel activation, or both^12–15^. Recent targeted therapies rescue CFTR cell surface localization (correctors) or channel function (potentiators), these therapies have been transformative for the health and lifespan of CF patients^14^.

As both cystic fibrosis and ADPKD are caused by variants in ion channels, we wondered whether pathogenic variants in PC1 can be similarly classified and addressed with therapeutics. Here, we developed assays that allow functional classification of ADPKD variants throughout PC1 to understand how human disease-causing variants disrupt polycystin localization and function.

## Results

### Contribution of missense variants to ADPKD

It remains challenging to assess the pathogenicity of missense variants in PC1, potentially limiting prior estimates of the fraction of ADPKD explained by missense variants. We examined the incidence of truncating (frameshift and stop-gain) and missense variants in PC1 in patients with and without a diagnosis of polycystic kidney disease (PKD) in the UK Biobank. The UK Biobank cohort includes over 400,000 exome-sequenced individuals in which genotype is linked to phenotype including diagnostic codes for PKD.

We observed that 19.6% of individuals with PKD carry a truncating variant in PC1, whereas 0.02% of controls have such mutations (**Figure 1A**). This difference between the prevalence of truncating variants in cases and controls indicates that ∼20% of PKD diagnoses in UK Biobank are attributable to truncating variants. Similarly, 15.4% of patients with PKD carry a rare (allele frequency < 2*10^−5^) missense variant, compared to 1.8% of controls who carry a rare missense variant (**Figure 1A**). Thus, missense variants in PC1 account for ∼14% of PKD-affected individuals the in UK Biobank. We note that this difference between the carrier frequency of rare missense variants in ADPKD cases and controls is robust to the choice of allele frequency (Supplementary Figure 1). These data suggest that PC1 missense variants account for nearly as many cases of PKD as PC1 truncating variants.

**Figure 1.**
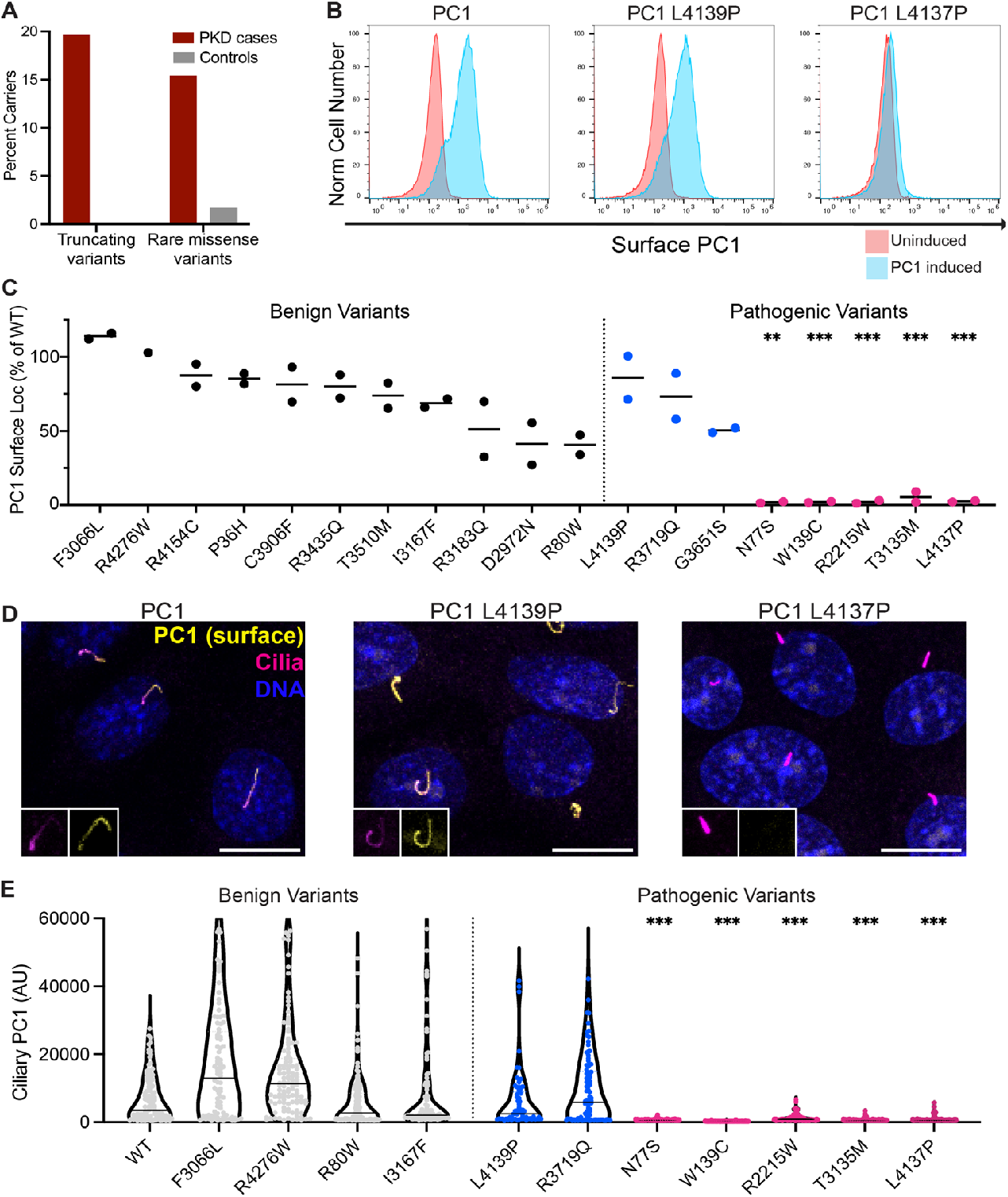
Many pathogenic ADPKD variants disrupt polycystin cell surface localization. A. Nontruncating and truncating variants in PC1 explain a similar proportion of polycystic kidney disease diagnoses. The fraction of individuals with a truncating variant or rare missense variant is shown for individuals with or without a diagnosis of polycystic kidney disease in the UK Biobank. B. Cell surface localization of wildtype PC1 or PC1 with pathogenic variants in nonciliated HEK293 cells measured by flow cytometry. PC1 expression is doxycycline inducible, cells cultured without doxycycline serve as a control. C. Quantitation of cell surface localization of PC1 by flow cytometry for benign and pathogenic variants in HEK293 cells. Each point represents an independent experiment; localization is plotted as a fraction of wild-type PC1 expression from the same experiment. Significance was determined with an ANOVA with Dunnett’s multiple comparison test, ** p<0.01,*** p<0.001. D. Cell surface localization of wildtype PC1 or PC1 with pathogenic variants in ciliated IMCD3 cells visualized by immunofluorescence. ARL13B is stained to indicate primary cilia. Scale bar, 10uM. E. Ciliary membrane PC1 variant expression in IMCD3 cells quantified by immunofluorescence is shown in violin plots with the median indicated. Significance was determined with an ANOVA on ranks with a one-sided Dunn’s multiple comparison test, *** p<0.01.

### The impact of pathogenic Polycystin-1 variants on cell surface localization

Evaluating pathogenicity of individual missense variants in PC1 remains challenging due to the small fraction of cases caused by each variant (allelic heterogeneity) and the difficulty in functionally assessing these variants. We therefore combined data from ClinVar^16^, the Mayo ADPKD variant database^2^, allele frequency, and published literature to identify probable benign and pathogenic variants throughout PC1 for further study (see methods). We then generated PC1 expression constructs with single amino acid pathogenic and benign variants (Supplemental Table 1).

We tested whether pathogenic missense variants in PC1 impair cell surface localization using our previously published PC1 expression construct^17^. This construct includes a modified PC1 in which the endogenous signal peptide was replaced with the 12 amino acid IgG leader sequence followed by an HA tag^17^. When co-expressed with PC2, this construct allowed us to selectively visualize PC1-containing polycystin channels at the ciliary membrane of kidney-derived IMCD3 cells. In non-ciliated HEK-293 cells, these expression constructs have allowed us to measure polycystin channel function at the plasma membrane^17^.

We generated stable HEK-293 cell lines co-expressing PC1 missense variants together with PC2 and quantified plasma membrane localization of the polycystin complex using the HA-tag on PC1 by flow cytometry. As PC1 and PC2 are expressed under the control of a doxy-cycline-inducible promoter, cells cultured in the absence of doxycline served as controls. Flow cytometry detected robust cell surface localization of wildtype PC1 (**Figure 1B**). Similarly, the pathogenic PC1 variant L4139P^10^ localized to the cell surface equivalently to wild-type PC1. In contrast, the pathogenic variant L4137P^10^ exhibited markedly reduced cell surface levels. Thus, similar PC1 variants separated by a single residue can have markedly different effects on polycystin localization.

For a more global view of the effect of genetic variants on PC1, we measured cell surface localization of 11 benign and 8 pathogenic PC1 variants (**Figure 1C**). All benign missense variants localized robustly at the cell membrane (**Figure 1C**). In contrast, 5 out of 8 tested pathogenic variants (N77S, W139C, R2215W, T3135M, L4137P) attenuated localization to the plasma membrane (**Figure 1C**, magenta dots). We conclude that impaired surface localization of the polycystin complex may be a common underlying mechanism for ADPKD.

Endogenous polycystins localize to the primary cilium of kidney epithelial cells. Loss of cilia or ciliary components, like loss of polycystins, causes kidney cysts suggesting that polycystins function at the primary cilium^8,9^. We therefore asked whether pathogenic variants in PC1 affect ciliary polycystin localization. Consistent with plasma membrane localization in HEK-293 cells, all benign PC1 variants robustly localized to the primary cilia of IMCD3 cells (**Figure 1D,E**). Moreover, the 5 pathogenic PC1 variants that compromised plasma membrane localization in HEK-293 cells also failed to localize to primary cilia of IMCD3 cells (**Figure 1D,E**, magenta dots). These pathogenic PC1 variants remain intracellular (Supplementary Figure 2). These results indicate that a subset of pathogenic ADPKD variants prevent polycystin localization to cilia.

### The impact of pathogenic Polycystin-1 variants on polycystin channel function

A subset of pathogenic ADPKD variants still efficiently localized to the plasma and ciliary membrane equivalently to benign variants (G3651S, R3719Q, L4139P; **Figure 1C** and **1E**, blue dots). This suggests that these variants might functionally compromise PC1 through a distinct mechanism such as impaired channel function. To test this hypothesis, we co-expressed pathogenic PC1 variants with a gain-of-function mutation in PC2 (F604P, PC2GOF) in HEK-293 cells, and measured polycystin currents in whole cell recordings (**Figure 2A,B**)^17,18^. As previously observed, cells coexpressing wildtype PC1 with PC2GOF generated robust outward-ly-rectifying currents (71.00±6.60 pA/pF, n=21, **Figure 2C,D**). In contrast, cells coexpressing PC2GOF with PC1 G3651S (7.27± 0.99 pA/pF, n=11), PC1 R3719Q (8.65±1.07 pA/pF, n=12), and PC1 L4139P (14.67±2.13 pA/pF, n=13) exhibited dramatically decreased channel activity, indistinguishable from background currents (**Figure 2C,D**). Together, these experiments suggest that some pathogenic PC1 variants that do not affect PC1 localization impair channel activity. We propose that there are at least two functional groups of nontruncating pathogenic PC1 variants: those which disrupt polycystin localization and those which disrupt channel activity.

**Figure 2.**
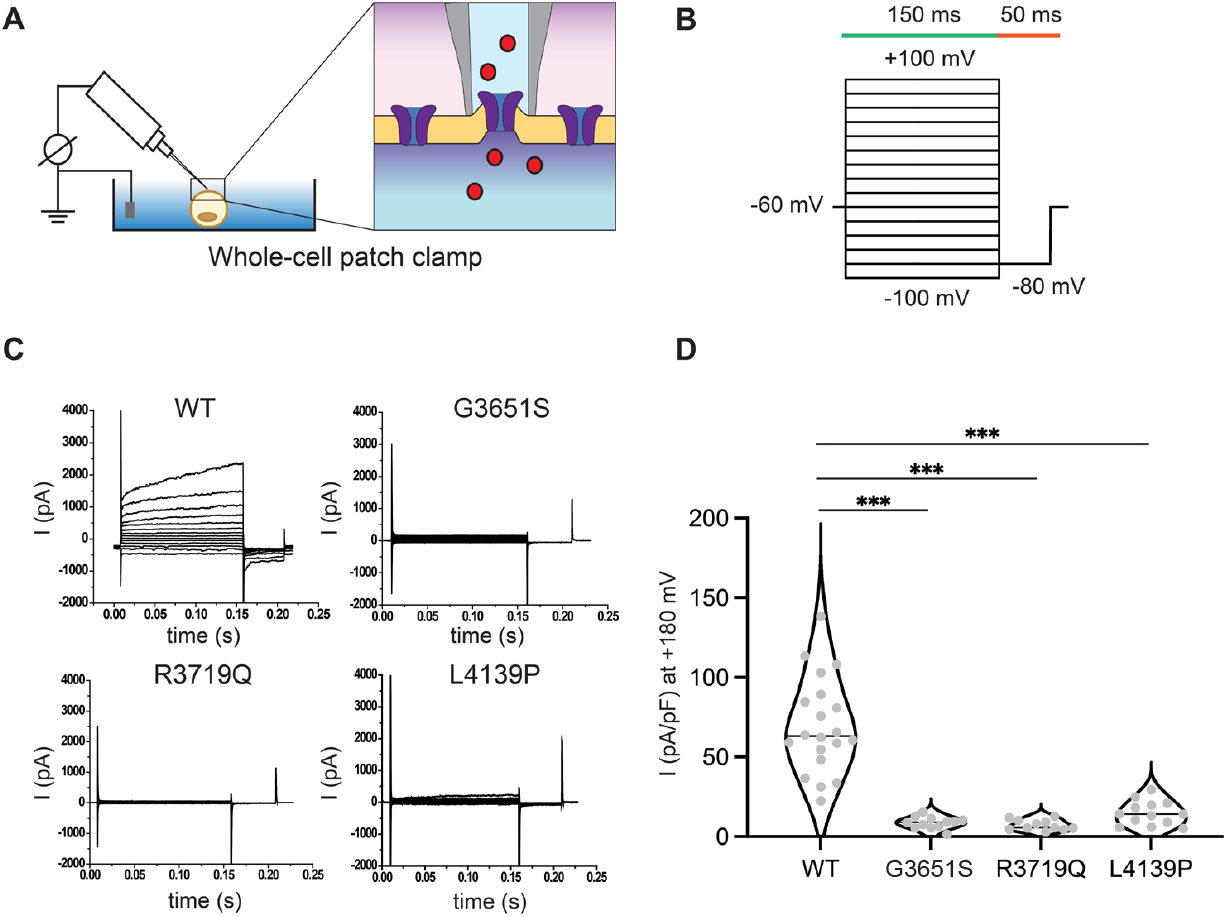
Pathogenic variants with normal surface expression disrupt polycystin channel function. A. Whole-cell patch clamp was used to measure polycystin channel function in the plasma membrane of HEK-293 cells. B. Step pulse protocol applied for the whole cell patch clamp. The voltage step was given from −100 mV to +180 mV in −20 mv increments. Each pulse was given for 150 ms and −80 mV tail pulse was followed for 50 ms. Holding potential was maintained at −60 mV. C. Representative whole cell patch clamp recordings for WT PC1, G3651 PC1, R3719Q PC1, and L4137P PC1. D. Mean current density (pA/pF) of wildtype and pathogenic variants. The current density was obtained at 140 ms after +180 mV step was applied. Wildtype n=20, G3651S n=12, R3719Q n=11, and L4139P n=13. Significance was determined with an ANOVA with Dunnet’s multiple comparison test, *** p<0.001.

### Classification of Variants of Uncertain Significance in Polycystin-1

There are no prevalent alleles in ADPKD; this allelic heterogeneity makes variant interpretation challenging and limits the utility of genetic testing for patients with missense PC1 variants. We therefore asked whether we could functionally classify PC1 missense variants of uncertain significance (VUS) carried by individuals with ADPKD (**Figure 3A,B**).

**Figure 3.**
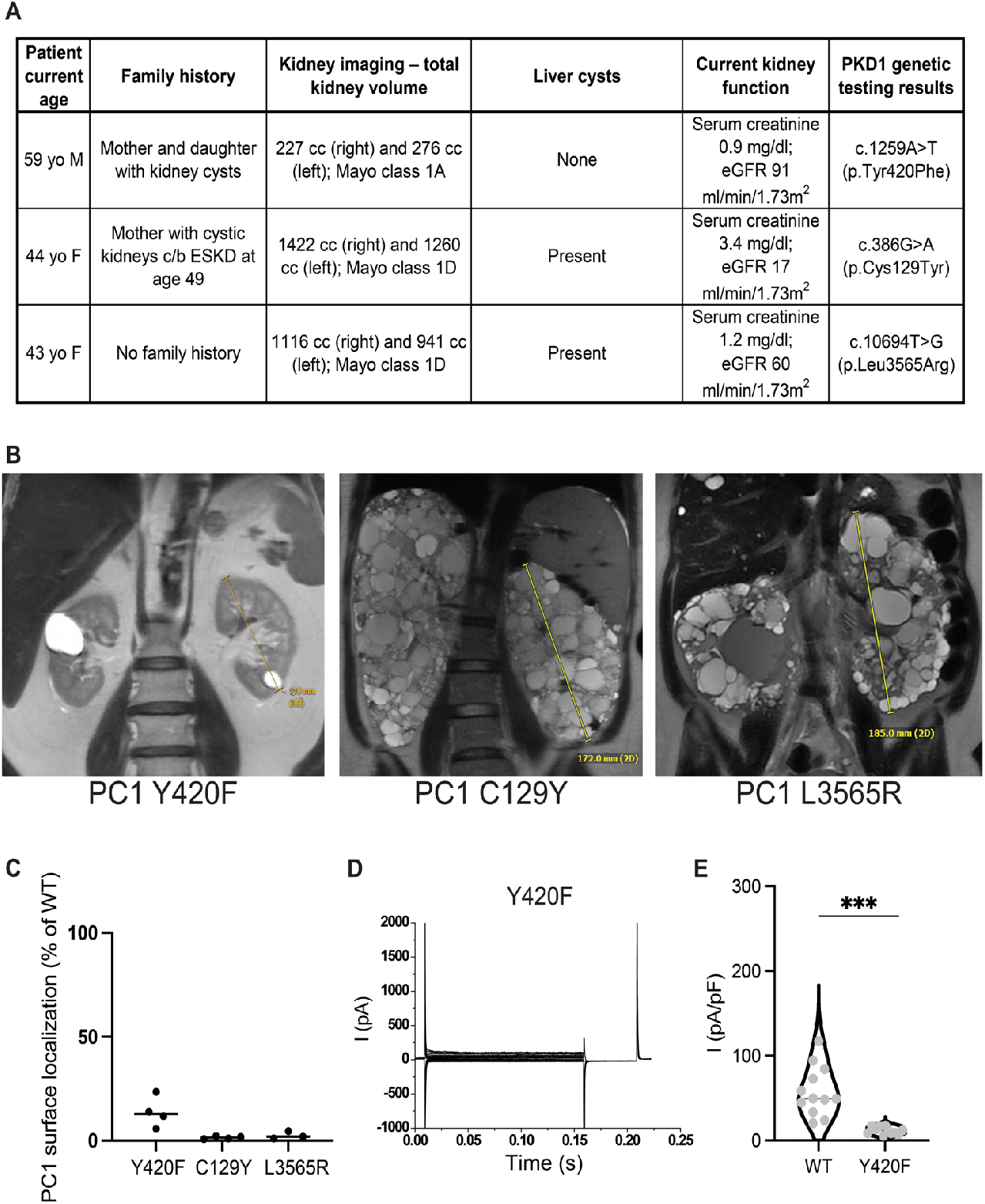
Classifying the effects of variants of uncertain significance on polycystin surface localization and channel function. A. Clinical and genetic characteristics of patients with ADPKD and VUSs in PC1. B. Abdominal MRIs from patients with ADPKD and VUSs in PC1. C. Reduced cell surface localization of VUSs in PC1 identified in patients with ADPKD. Cell surface localization was quantified by flow cytometry in HEK-293 cells and is plotted relative to wild-type PC1 expression from the same experiment. Each point represents an independent experiment. D. Representative whole cell patch clamp recording for Y420F PC1. E. Mean current density (pA/pF) of WT PC1 and PC1 Y420F. The current density was obtained at 140 ms after +180 mV step pulse was applied. WT n= 12, Y420F n=11. Significance was determined with an unpaired t-test, *** p<0.001.

We generated three HEK-293 cell lines, each co-expressing one of the three PC1 VUS together with PC2 and quantified cell surface localization. PC1 C129Y and L3565R did not localize to the cell surface. In contrast, PC1 Y420F exhibited decreased but measurable cell surface localization (**Figure 3C**). We also measured channel function of PC1 Y420F coexpressed with PC-2GOF in whole cell recordings. Cells expressing PC1 Y420F/PC2GOF had dramatically impaired channel activity compared to cells expressing wild-type PC1/PC-2GOF (**Figure 3D,E**). These results suggest that combined surface expression and ion channel function assays hold promise for classifying pathogenicity of PC1 variants.

### Restoring ciliary localization of Polycystin-1 pathogenic variants

Since many ADPKD-causing variants impair polycystin surface localization, pharmacological interventions to increase surface localization may be therapeutically beneficial. The CF-causing mutation CFTR F508del, poorly localizes to the plasma membrane; however decreased temperature increases F508del surface localization^19^. Therefore, we investigated whether decreased temperature increases polycystin cell surface localization. More specifically, we cultured IMCD3 cells expressing PC1 variants with PC2 at 37°C and 32°C and assessed PC1 ciliary and cell surface localization. Ciliary localization of wildtype PC1 at 37°C and 32°C was equivalent (**Figure 4A,B**). In contrast, ciliary localization of PC1 L4137P and R2215W, two variants that compromise ciliary localization at 37°C, was augmented after 24h at 32°C (**Figure 4A,B**). We also tested the ability of lowering temperature to restore plasma membrane localization of PC1 variants with surface localization defects. In HEK-293 cells expressing PC1 variants together with PC2, we quantified PC1 cell surface localization after 24h at 37°C or 32°C. Cell surface localization of PC1 N77S and PC1 W139C was unaffected by temperature (**Figure 4C**). In contrast, cell surface localization of PC1 R2215W and T3135M increased at 32°C (**Figure 4C**). Thus, a subset of PC1 variants exhibit temperature-sensitive cell surface localization.

**Figure 4.**
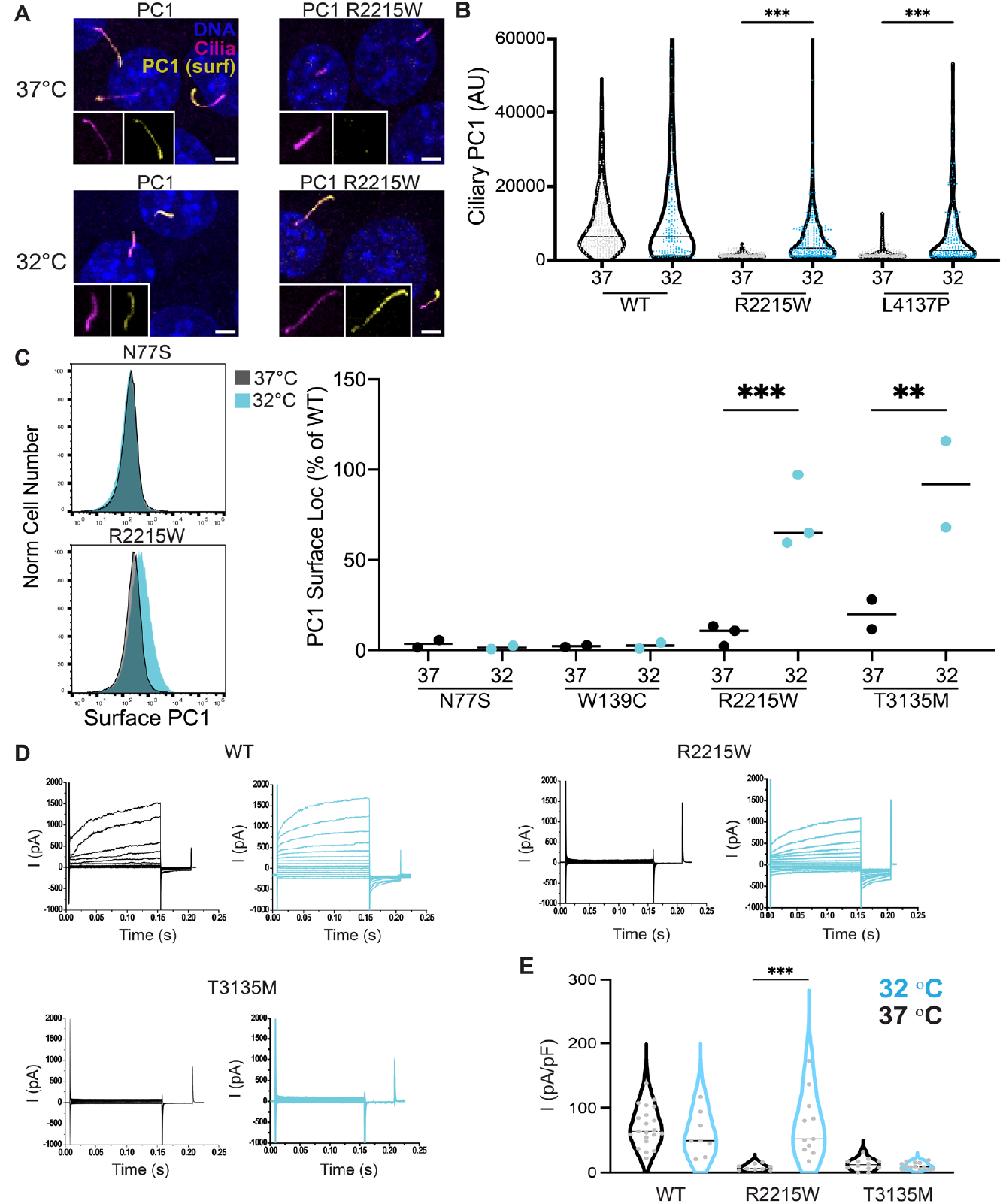
The effect of temperature on localization of pathogenic PC1. A. Ciliary expression of wildtype PC1 and pathogenic R2215W PC1 in IMCD3 cells cultured at 37°C and 32°C Scale bar, 2.5uM. B. Increased ciliary localization of pathogenic L4137P PC1 and R2215W PC1 in IMCD3 cells cultured at 32°C. Violin plots are shown with the median represented by a black line. Significance was determined with an ANOVA on ranks with a Dunn’s multiple comparison test. ** indicates p<0.01. C. Flow cytometry of cell surface expression of PC1 pathogenic variants in HEK293 cells cultured at 37°C or 32°C (left) and quantified relative to wildtype PC1 measured in the same experiment (right). Each point represents an independent experiment. Significance was determined with an ANOVA with the Šidák multiple comparison test, ** p<0.01, *** p<0.001. D. Representative whole cell patch clamp recordings for cells expressing the indicated PC1 variant cultured at 37°C (black) or 32°C (blue). E. Mean current density of cells expressing the indicated PC1 variant cultured at 37°C or 32°C shown as violin plots with median indicated. WT 37°C n=21, 32°C n=8, R2215W 37°C n=9, 32°C n=11, T3135M 37°C n=12 32°C n=13. Significance was determined with an ANOVA with the Šidák multiple comparison test, *** p<0.001.

We investigated whether polycystin variants rescued by low temperature retained channel activity. We measured polycystin channel activity in whole cell patch-clamp recordings of HEK-293 cultured at either 32°C or 37°C. Whereas lowering temperature restored cell surface localization of PC1 T3135M/ PC2GOF, it did not restore current densities (37°C: 10.21±1.38 pA/pF, n=13; 32°C:12.48±2.61 pA/pF, n=12), indicating that the T3135M variant disrupts both surface localization and channel function (**Figure 4D,E**). In contrast, PC1 R2215W/PC2GOF generated robust currents when surface localization was restored by culture at 32°C (37°C: 7.74±1.63 pA/pF, n=10; 32°C: 73.06±14.55 pA/pF, n=11), indicating that the R2215W variant specifically disrupts surface localization and does not compromise channel function. These experiments indicate that restoring surface localization of a subset of PC1 pathogenic variants is sufficient to rescue channel function.

To determine whether PC1 surface localization may also be promoted by small molecules, we tested three correctors identified for their capacity to promote cell surface localization of CFTR F508del. All three compounds (C2, C3 and C18; Cystic Fibrosis Foundation) had negligible effects on the ciliary localization of wild-type PC1 (Supplementary Figure 3). In contrast, one small molecule, C3, also known as VRT-325, increased ciliary localization of PC1 R2215W and L4317P (**Figure 5A,B**, Supplementary Figure 3). Thus, small molecules can increase ciliary localization of some ADPKD-causing PC1 variants.

**Figure 5.**
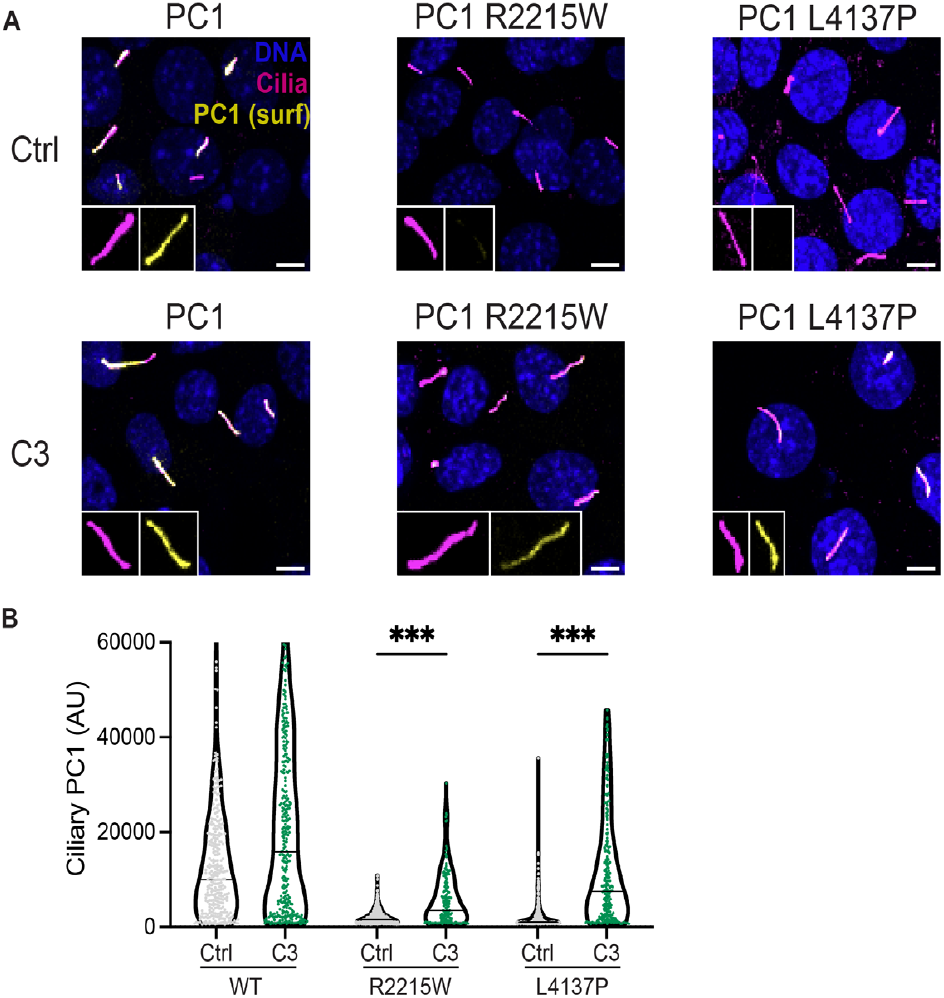
Small molecule rescue of PC1 ciliary localization. Representative cell surface localization of the indicated PC1 variant in the presence or absence of the small molecule C3. Surface PC1 expression and the ciliary marker ARL13B were visualized by immunofluorescence. Scale bar, 5uM. Ciliary membrane PC1 variant localization in IMCD3 cells shown as violin plots with the median indicated. Significance was determined with an ANOVA on ranks with a Dunn’s multiple comparison test, *** p<0.001.

## Discussion

### Nontruncating variants in PC1 account for ∼40% of ADPKD attributable to PC1

We measured the prevalence of both PC1 truncating variants and rare missense variants in individuals with polycystic kidney disease and controls in the UK Biobank. Comparing the prevalence of these two types of variants suggests that missense variants contribute to ∼40% of ADPKD caused by PC1 variants. This proportion agrees with prior studies in ADPKD-specific cohorts, which used completely different strategies to classify variants and patients and did not include control populations^3,4,10^. The total fraction of PKD explained by PC1 variants in the UK Biobank (∼35%) is less than is seen in ADPKD clinical cohorts (∼80%), likely because some individuals with a PKD diagnostic code do not have ADPKD. However, this is unlikely to affect the estimate of the ratio between patients with truncating and missense variants. In addition, our analysis revealed that 1.8% of healthy individuals in the UK Biobank have a rare PC1 variant (allele frequency < 2×10^−5^) and thus in the human population rare benign missense variants in PC1 vastly exceed the number of rare pathogenic variants (by ∼200:1). Thus, missense variants contribute to a large fraction of ADPKD, and classification of these variants to improve genetic testing and inform potential therapeutic strategies is sorely needed.

### Pathogenic ADPKD variants disrupt polycystin surface localization or channel function

Interrogating polycystin localization and channel activity allowed us to characterize molecular mechanisms that may underlie ADPKD. In a randomly chosen group of pathogenic PC1 variants, including eight pathogenic variants and three VUSs in ADPKD-patients, seven variants impaired PC1 cell surface localization, three affected channel function, and one disrupted both. We propose that ADPKD pathogenic missense variants fall into two major groups, based on cellular phenotypes. Some disrupt PC1 surface localization and others impair channel function. These groups are not mutually exclusive as shown for PC1 Y420F. We currently only have a limited understanding how variants in PC1 affect cell surface localization of the PC1/PC2 heterotetramer. As pathogenic variants disrupted both ciliary localization and plasma membrane localization in non-ciliated cells, most are likely to disrupt processes fundamental to polycystin biogenesis or stability rather than specifically disrupting ciliary trafficking. Biogenesis of most transmembrane proteins requires translation at the rough ER, folding, modification in the Golgi and vesicular trafficking^20^. In addition, PC1 biogenesis requires glycosylation^21–23^, autoproteolytic cleavage^24^, and assembly with PC2^25^. As reduced temperatures improved surface localization of some PC1 variants, these variants may affect protein folding^19,26,27^. Determining whether non-temperature sensitive variants predominantly affect protein folding, protein-protein interactions essential for biogenesis, or protein stability may help further stratify these variants for future therapeutic development.

A second group of pathogenic variants localized normally to the ciliary membrane but displayed attenuated channel activity. It is unclear how these variants affect channel activation. The R3719Q variant resides in the extracellular loop between TM6 and TM7 of PC1 called the TOP domain. The TOP domain is also present in PC2 and pathogenic variants in the PC2 TOP domain exhibit a change in voltage-dependent channel activation^28^. Future studies will be needed to address how G3651S or L4139P affect channel activation and whether channel activity in these and other pathogenic variants can be restored with small molecules.

Prior work has indicated that the polycystins can function as ion channels^29^, atypical GPCRs^30–32^, or that their cleavage products can act as signaling mediators^33,34^. The relevance of these functions for human ADPKD remains unclear. Our work supports the hypothesis that polycystin ion channel function is critical to ADPKD pathogenesis in agreement with a previous report^28^. In the future, testing surface localization and channel function defects of larger numbers of variants may identify additional functional categories of PC1 variants in ADPKD pathogenesis.

### A small molecule restores ciliary polycystin localization and suggests an approach for ADPKD therapy

The disruption of polycystin surface localization and channel function by ADPKD pathogenic variants is reminiscent of cystic fibrosis. Recently developed small molecule therapeutics for CF either: 1) promote surface localization of pathogenic CFTR variants (called correctors) or 2) increase channel function (called potentiators). Today the combination of correctors and potentiators treat ∼90% of all CF patients, converting the prior “death sentence” of CF into a manageable disease^14^. Our work suggests that similarly correctors and potentiators for polycystins hold promise for ADPKD. We found that a small molecule C3, improved ciliary localization of some pathogenic ADPKD variants. Given that each ADPKD variant accounts for a small fraction of cases, identifying therapies that are broadly useful over functional classes of pathogenic variants is likely to be important.

In summary, our results reveal an approach to classify AD-PKD-causing missense variants and suggests the potential of therapies that promote polycystin surface localization and channel function.

## Acknowledgements

We thank the patients described in this article for their generosity of spirit in sharing their stories for the greater PKD community. This work was supported by NIH K99DK131361 (KH), NIH T32DK007219 and the UCSF Physician Scientist Scholars Program (GL), and NIH R01DK127277 (MD). We thank Feng Qian and the Polycystic Kidney Disease Research Resource Consortium (U54DK126114) for providing IMCD3 cells with CRISPR ablated PC1, the Cystic Fibrosis Foundation for providing CFTR correctors, and the participants and leadership of the UK Biobank.

## Methods

### UK Biobank Analysis

UK Biobank subjects with at least one ICD-10 code and exome data (415,848) were included in the analysis. The ICD-10 code Q61.2 (Polycystic Kidney Disease, Adult) was used to identify PKD cases. Genotypes were ascertained using the population level exome OQFE data (final release) in the pVCF format. Variants were initially normalized (with ‘bcftools norm’), followed by only retaining variants for which at least 90% of the samples had a non-missing genotype and at least 10 reads. Missense, stop gain, and frameshift variant annotations were determined using Ensembl Variant Effect Predictor^35^. Frameshift or stop gained variants, were jointly labeled as truncating variants. For Supplementary Figure 3, each subject was assigned an AF based on the rarest missense variant they had in PKD1, or an AF of 1 if they had no missense variants in PKD1. This work was carried out under UK Biobank Application 78378.

### Variant Classification

We defined likely benign variants as those with allele frequency greater than 2×10^−5^, reported homozygosity in humans—as loss of PC1 is embryonic lethal^36^, and classified as likely benign or benign in the Mayo PKD database or Clinvar. We defined likely pathogenic variants as those with allele frequencies less than 5×10^−6^, reports of pathogenicity in the literature, and classification of likely pathogenic or pathogenic in the Mayo PKD database or Clinvar. Variants of uncertain significance (VUS) were classified based on clinical genetic testing from a reference laboratory (Natera).

### Patients

Patients with VUS in PKD1 provided individual consent for participation in the description of their clinical histories and genetic variants. Human subjects research at the UCSF PKD Center of Excellence is performed under IRB #14-15601.

### Molecular biology

κHA-PC1 has been described previously^17^. In short, the endogenous signal peptide in PC1 was replaced with the 12aa IgG leader sequence followed by an HA tag. All missense mutations in PC1 were introduced using PCR amplification and Gibson assembly (NEB) using several unique restriction sites in PC1. PC1 variants were introduced into a plasmid (Addgene #104454) containing a hygromycin selection cassette. PC2 was also ligated into the same doxycycline inducible PiggyBac expression plasmid as PC1 but with a puromycin selection cassette instead. All DNA sequences were confirmed by whole plasmid sequencing (Primordium Labs).

### Antibodies

Rat anti-hemagglutinin (HA), Roche (11867423001); Rabbit anti-hemagglutinin (HA) (C29F4), Cell Signaling Technology (3724); Mouse anti-Arl13b (N295B/66), Antibodies Inc.

### Immunocytochemistry and confocal microscopy

Cells were fixed with 4% formaldehyde, permeabilized with 0.2% Triton X-100, and blocked by 2% FBS, 2% BSA and 0.2% fish gelatin in PBS. Cells were labeled with the indicated antibody and secondary goat anti-rabbit, anti-rat or anti-mouse fluorescently labeled IgG (Thermo Fisher) and Hoechst 33342 (Thermo Fisher). Confocal images were obtained using a Zeiss LSM 800 laser scanning confocal microscope equipped with a 63x/1.4 oil immersion objective. Images were further processed using ImageJ (NIH).

### Cell culture and quantification of ciliary and plasma membrane PC1 and PC2

Surface localization of PC1/PC2 complexes was measured by quantifying amount of anti-HA bound to the plasma membrane. First we isolated isogenic clones of HEK293 or IMCD3 cells with CRISPR ablated endogenous PKD1 expression (gift of F.Qian^17^) only expressing PC2 under a doxycycline-inducible promoter. PC1 variants were introduced with the constructs described above and stable cell lines were generated using hygromycin selection. This approach ensured comparable PC2 expression levels between individual PC1 mutant cell lines. Polycystin complex expression was induced in HEK-293 cell lines with 2 ug/ ml doxycycline in complete medium for 48h. For experiments of the effect of temperature on surface localization, the last 24h of induction was performed at the indicated temperature (37°C or 32°C). Cells were detached with Versene (Thermo Fisher) and incubated on ice in blocking buffer (2% FBS, 2% BSA and 0.2% fish gelatin in PBS) for 10 minutes. After centrifugation cell pellets were resuspended in blocking buffer with rabbit anti-HA diluted 1:100. After incubation on ice for 20 min, cells were washed twice with cold PBS and incubated for 20 min with 488-conjugated goat anti-rabbit antibodies. Cells were washed again 2x with ice-cold PBS. Sytox Blue (Thermo Fisher) was added at 1:1000 5 minutes prior to sample acquisition on an Attune Nxt flow cytometer. Flow cytometry analysis was performed using FlowJo (v10.8.2). PC1 surface localization was quantified as the fraction of live cells with PC1 surface localization greater than an uninduced control.

To quantify the amount of polycystin complex in primary cilia, IMCD3 cell lines were first incubated in Opti-MEM (Thermo Fisher) for 24h to initiate ciliogenesis. Cells were further incubated for 48h with 0.2 ug/ ml doxycycline in Opti-MEM. For surface staining adherent live cells were incubated with L-15 medium containing rabbit anti-HA antibody (1:100) for 25 min at room temperature to avoid internalization of antibodies. Cells were washed twice with L-15 medium and fixed with 4% PFA and permeabilized with 0.2% Triton X-100. 8. Surface HA fluorescence was quantified from Z-projections in regions of interest defined by Arl-13b staining (cilia marker) using ImageJ. This approach allowed an unbiased quantification of membrane inserted HA.

For experiments using small molecule correctors, IMCD3 cells were incubated in Opti-MEM for 24h to initiate ciliogenesis, after which they were incubated for 48h with 0.2 ug/ml doxycycline and 5uM of the small molecule corrector in Opti-MEM. The following small molecules were used: C2 (VRT-640): 2-{1-[4-(4-Chloro-benzensulfonyl)-piperazin-1-yl]-ethyl}-4-piperidin-1-yl-quinazoline, C3 (VRT-325): 4-Cyclohexyloxy-2-{1-[4-(4-methoxybenzensulfonyl)-piperazin-1-yl]-ethyl}-quinazoline, C18 (VRT-534): 1-(benzo [d][1,3]dioxol-5-yl)-N-(5-((S)-(2-chlorophenyl)((R)-3-hydroxypyrrolidin-1-yl)methyl)thiazol-2-yl)cyclopropanecarboxamide.

### Electrophysiology

We performed a whole cell configuration patch clamp using multiclamp 200B (Axon instruments) and digidata 1324A (Axon instruments). The amplitudes from the polycystin complex and other variants were recorded using pClamp software (Axon instruments). Whole cell configuration patch clamp data set was filtered at 1kHz and sampled at 10 kHz. The voltage step pulse ranged from −100 mV to 180 mV in 20 mV increments during 150 ms followed by −80 mV tail pulse during 50 ms. The holding potential was given at −60 mV for the recordings. We pulled the glass pipettes using P-100 micropipette puller (Sutter instrument). The resistance of pipettes for the whole cell recordings was within 6-8 MΩ range. The tip of pipette was further polished using a Narishige MF-830 microforge. For patch clamp experiments, we used an extracellular solution consisting of (mM): 145 Na-gluconate, 5 KCl, 2 CaCl2, 1 MgCl2, 10 HEPES and adjusted to pH 7.4 using NaOH. We used an intracellular solution consisting of (mM): 90 NaMES, 10 NaCl, 2 MgCl2, 10 HEPES, 5 EGTA, 100 nM free calcium adjusted by CaCl2 and adjusted to pH 7.4 using NaOH.

We used Clampfit10.6 (Axon Instruments/Molecular devices), Origin8 (Originlab), and Prism10.0 (Graphpad) to analyze data from the whole cell configuration patch clamp. Data are shown as mean ± s.e.m., and n represents the number of tested cells.

## Supplementary Information

**Supplementary Figure 1:** Prevalence of missense variants in PKD cases and controls by allele frequency.

**Supplementary Figure 2:** Total and cell surface expression of PC1.

**Supplementary Figure 3:** Effect of small molecule correctors on ciliary PC1 expression.

## References

1. Chapman, A. B. et al. Autosomal-dominant polycystic kidney disease (ADPKD): executive summary from a Kidney Disease: Improving Global Outcomes (KDIGO) Controversies Conference. Kidney International 88, 17– 27 (2015).

2. ADPKD Variant Database. https://pkdb.mayo.edu/vari-ants.

3. Cornec-Le Gall, E. et al. Type of PKD1 Mutation Influences Renal Outcome in ADPKD. J Am Soc Nephrol 24, 1006–1013 (2013).

4. Mantovani, V. et al. Gene Panel Analysis in a Large Cohort of Patients With Autosomal Dominant Polycystic Kidney Disease Allows the Identification of 80 Potentially Causative Novel Variants and the Characterization of a Complex Genetic Architecture in a Subset of Families. Front Genet 11, 464 (2020).

5. Mochizuki, T. et al. PKD2, a Gene for Polycystic Kidney Disease That Encodes an Integral Membrane Protein. Science 272, 1339–1342 (1996).

6. The European Polycystic Kidney Disease Consortium. The polycystic kidney disease 1 gene encodes a 14 kb transcript and lies within a duplicated region on chromosome 16. Cell 77, 881–894 (1994).

7. Su, Q. et al. Structure of the human PKD1-PKD2 complex. Science 361, eaat9819 (2018).

8. Yoder, B. K., Hou, X. & Guay-Woodford, L. M. The polycystic kidney disease proteins, polycystin-1, polycystin-2, polaris, and cystin, are co-localized in renal cilia. J. Am. Soc. Nephrol. 13, 2508–2516 (2002).

9. Pazour, G. J., San Agustin, J. T., Follit, J. A., Rosenbaum, J. L. & Witman, G. B. Polycystin-2 localizes to kidney cilia and the ciliary level is elevated in orpk mice with polycystic kidney disease. Curr Biol 12, R378–380 (2002).

10. Rossetti, S. et al. Comprehensive molecular diagnostics in autosomal dominant polycystic kidney disease. J Am Soc Nephrol 18, 2143–2160 (2007).

11. Riordan, J. R. et al. Identification of the cystic fibrosis gene: cloning and characterization of complementary DNA. Science 245, 1066–1073 (1989).

12. Sheppard, D. N. et al. Mutations in CFTR associated with mild-disease-form Cl-channels with altered pore properties. Nature 362, 160–164 (1993).

13. Li, C. et al. ATPase activity of the cystic fibrosis transmembrane conductance regulator. J Biol Chem 271, 28463–28468 (1996).

14. Grasemann, H. & Ratjen, F. Cystic Fibrosis. New England Journal of Medicine 389, 1693–1707 (2023).

15. Cheng, S. H. et al. Defective intracellular transport and processing of CFTR is the molecular basis of most cystic fibrosis. Cell 63, 827–834 (1990).

16. Landrum, M. J. et al. ClinVar: improving access to variant interpretations and supporting evidence. Nucleic Acids Res 46, D1062–D1067 (2018).

17. Ha, K. et al. The heteromeric PC-1/PC-2 polycystin complex is activated by the PC-1 N-terminus. eLife 9, e60684 (2020).

18. Arif Pavel, M. et al. Function and regulation of TRPP2 ion channel revealed by a gain-of-function mutant. Proc Natl Acad Sci U S A 113, E2363–2372 (2016).

19. Denning, G. M. et al. Processing of mutant cystic fibrosis transmembrane conductance regulator is temperature-sensitive. Nature 358, 761–764 (1992).

20. Hegde, R. S. & Keenan, R. J. The mechanisms of integral membrane protein biogenesis. Nat Rev Mol Cell Biol 23, 107–124 (2022).

21. Besse, W. et al. Isolated polycystic liver disease genes define effectors of polycystin-1 function. J Clin Invest 127, 1772–1785 (2017).

22. Besse, W. et al. ALG9 Mutation Carriers Develop Kidney and Liver Cysts. Journal of the American Society of Nephrology 30, 2091 (2019).

23. Porath, B. et al. Mutations in GANAB, Encoding the Glucosidase IIα Subunit Cause Autosomal-Dominant Polycystic Kidney and Liver Disease. The American Journal of Human Genetics 98, 1193–1207 (2016).

24. Qian, F. et al. Cleavage of polycystin-1 requires the receptor for egg jelly domain and is disrupted by human autosomal-dominant polycystic kidney disease 1-associated mutations. Proc Natl Acad Sci U S A 99, 16981–16986 (2002).

25. Gainullin, V. G., Hopp, K., Ward, C. J., Hommerding, J. & Harris, P. C. Polycystin-1 maturation requires polycystin-2 in a dose-dependent manner. J Clin Invest 125, 607–620 (2015).

26. Sturtevant, J. M., Yu, M. H., Haase-Pettingell, C. & King, J. Thermostability of temperature-sensitive folding mutants of the P22 tailspike protein. J Biol Chem 264, 10693–10698 (1989).

27. Brown, C. R., Hong-Brown, L. Q. & Welch, W. J. Correcting temperature-sensitive protein folding defects. https://www.jci.org/articles/view/119302/pdf (1997) doi:10.1172/JCI119302.

28. Vien, T. N., Wang, J., Ng, L. C. T., Cao, E. & DeCaen, P. G. Molecular dysregulation of ciliary polycystin-2 channels caused by variants in the TOP domain. Proceedings of the National Academy of Sciences 117, 10329–10338 (2020).

29. Hanaoka, K. et al. Co-assembly of polycystin-1 and -2 produces unique cation-permeable currents. Nature 408, 990–994 (2000).

30. Delmas, P. et al. Constitutive activation of G-proteins by polycystin-1 is antagonized by polycystin-2. J Biol Chem 277, 11276–11283 (2002).

31. Wu, Y. et al. Gα12 is required for renal cystogenesis induced by Pkd1 inactivation. J Cell Sci 129, 3675–3684 (2016).

32. Parnell, S. C. et al. The Polycystic Kidney Disease-1 Protein, Polycystin-1, Binds and Activates Heterotrimeric G-Proteinsin Vitro. Biochemical and Biophysical Research Communications 251, 625–631 (1998).

33. Onuchic, L. et al. The C-terminal tail of polycystin-1 suppresses cystic disease in a mitochondrial enzyme-dependent fashion. Nat Commun 14, 1790 (2023).

34. Woodward, O. M. et al. Identification of a polycystin-1 cleavage product, P100, that regulates store operated Ca entry through interactions with STIM1. PLoS One 5, e12305 (2010).

35. McLaren, W. et al. The Ensembl Variant Effect Predictor. Genome Biology 17, 122 (2016).

36. Lu, W. et al. Perinatal lethality with kidney and pancreas defects in mice with a targetted Pkd1 mutation. Nat Genet 17, 179–181 (1997).

